# PILOT-GM-VAE: Patient-Level Analysis of single cell Disease Atlas with Optimal Transport of Gaussian Mixture Variational Autoencoders

**DOI:** 10.1101/2025.04.10.648188

**Authors:** Mehdi Joodaki, Mina Shaigan, Samaneh Samiei, James Nagai, Tiago Maié, Christoph Kuppe, Ivan G. Costa

**Affiliations:** Institute for Computational Genomics, Joint Research Center for Computational Biomedicine, RWTH Aachen University Medical School, 52074 Aachen, Germany; Laboratory for Quantitative Cell Dynamics and Translational Systems Biology, RWTH Aachen University Medical School, 52074 Aachen, Germany

**Keywords:** Optimal Transport, Variational Autoencoder, Disease Progression, Mixture Models, scRNA-seq Disease Atlas, Patient-Level Modeling

## Abstract

**Motivation:** The analysis of single cell disease atlases represents a challenge due to the presence of batch effects, low quality of disease samples, and the multi-scale nature of the data, i.e., samples are described by different cell distributions. Because of these, few computational approaches are performing sample-level disease progression analysis so far.

**Results:** Here, we introduce Patient-Level Analysis with Optimal Transport based on Gaussian Mixture Variational Autoencoders (PILOT-GM-VAE). PILOT-GM-VAE explores the power of GM-VAE to estimate models describing complex single cell distributions through efficient optimal transport algorithms for estimating the distance between Gaussian Mixtures. Extensive benchmarking on several single cell disease atlases and competing approaches demonstrates the performance of PILOT-GM-VAE in sample-level clustering, sample-level trajectory inference, and batch correction tasks. Moreover, we performed a case study on a breast cancer disease atlas, where PILOT-GM-VAE highlighted cellular and molecular changes associated with breast cancer disease progression.

**Availability:** The software, code, and data for benchmarking are available at https://github.com/CostaLab/PILOT-GM-VAE/tree/main

## 1 Introduction

Single cell sequencing technologies have revolutionized our understanding of cellular diversity and function, enabling researchers to profile gene expression at an unprecedented resolution (Tang *et al*., 2009; Macosko *et al*., 2015), as shown by the recent efforts of the Human Cell Atlas measuring millions of cells in various organs (Yanai *et al*., 2024).

Although most of the atlases focuses on healthy tissues, there are increasing efforts to create disease single cell atlases, such as in lung diseases (Sikkema *et al*., 2023), covid (Ren *et al*., 2021), myocardial infarction (Kuppe *et al*., 2022), and kidney diseases (Lake *et al*., 2021), among others. The computational analysis of disease atlases is challenging due to the presence of experimental artifacts such as batch effects (Luecken *et al*., 2022), which are potentially worsened by the inherently lower quality of disease tissues compared to healthy tissue. Another challenge arises from the multi-scale nature of this data, i.e., every sample/patient is represented by a distribution of cells, which are in turn characterized by their genes. Such multi-scale data (samples x cells x genes) ^1^ cannot be straightforwardly analyzed by “standard” machine learning methods, which usually require tabular data (cells **x** genes or samples **x** genes).

Recently, a few computational approaches have been proposed for modeling a single cell experiment of a sample (or patient) as a distribution of cells. First, PhEMD (Phenotypic Earth Mover’s Distance; Chen *et al*. (2020)) employed Earth Mover’s Distance to measure distances between single cell samples based on diffusion-based clustering. This method was effective in assessing drug responses in cell lines. Yet, it relies on diffusion maps and pseudotime estimates that assume a continuous trajectory of cellular states, which do not necessarily exist in organs with heterogeneous cell populations. We have recently proposed PILOT (Patient-Level Analysis with Optimal Transport; (Joodaki *et al*., 2024)), which models samples as discrete distributions of cell types. PILOT uses optimal transport-based Wasserstein Distances(Peyré *et al*., 2019) to measure the distance between samples. A benchmarking analysis of eight disease scRNA-seq atlases indicated that PILOT had higher performance in clustering and trajectory inference problems when compared to PhEMD or pseudo-bulk patient’s single cell libraries. PILOT also includes the use of non-linear models for interpretation of results, i.e., associated cell types and genes related to the trajectory.

Later, Quantized Optimal Transport (QOT) was proposed, building upon the PILOT framework but using a mixture of Gaussian distributions to model the distribution of a particular cell type (Wang *et al*., 2025). As with PILOT, this method requires a previous annotation of cells to cell types. GloScope (Wang *et al*., 2024) estimates Gaussian Mixture Models (GMMs) independently for each sample to model cell distributions, while it uses a symmetric version of the Kullback-Leibler (KL) divergence to estimate a distance between distributions representing samples. Note also that GloScope was evaluated with limited benchmarking with few data sets and without a comprehensive comparison to all existing competing approaches.

## 2 Approach

In this work, we introduce a PatIent Level Optimal Transport approach with Gaussian Mixture Variational AutoEncoders (PILOT-GM-VAE). It uses VAE to estimate Gaussian mixture models representing the distribution of a single cell disease atlas. Next, PILOT-GM-VAE explores the fact that the Wasserstein Distance between Gaussian Mixtures (GMs) can be efficiently estimated (Chen *et al*., 2018) by using an analytical solution to transport two Gaussian distributions (Takatsu, 2011).

In detail, for a given integrated and dimensionally reduced single cell disease data, PILOT-GM-VAE uses an unsupervised version of the Variational Autoencoder (GM-VAE) proposed in Kingma *et al*. (2014) to estimate Gaussian Mixture models (Fig.1A). ^2^ Next, the GM-VAE latent variables are used to obtain a GM representing the distributions of cells in a given sample. The sample-specific GM models are used for estimation of the Wasserstein Distance between samples (Chen *et al*., 2018) (Fig.1B). The distances can be used for downstream analysis, such as sample-level clustering or sample-level disease progression, as well as to associate cellular states and molecular features (genes) associated with clusters or trajectories as before (Joodaki *et al*., 2024) (Fig.1C).

**Fig. 1:**
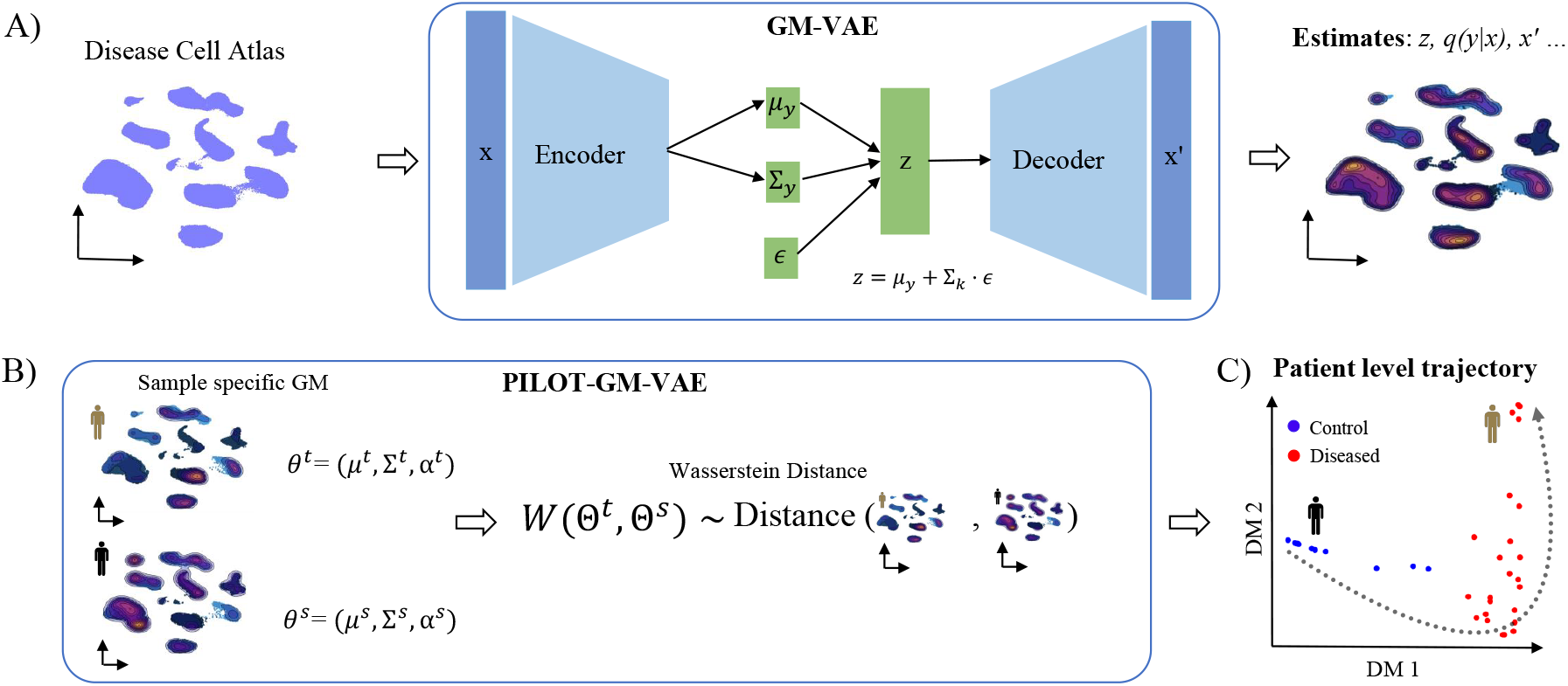
Schematic of the PILOT-GM-VAE framework. (A) PILOT-GM-VAE receives as input data an integrated scRNA-seq data matrix *x* from a disease atlas. Then, it uses a GM-VAE to encode *x* into a latent variable z conditional on the discrete variable y. The latter represents the component assignment in a mixture model. The GM-VAE outputs an estimated data x^0^ and posterior estimates on the association of data points to components *q*(*y*|*x*). (B) PILOT-GM-VAE explores the *q*(*y*|*x*) as estimated by the GM-VAE sample-specific GMs (Θ^*t*^ and Θ^*s*^). These are provided as input to the Wasserstein Distance based on the Bures-Wasserstein formulation. (C) The Wasserstein Distance between samples is used for downstream analysis, such as patient-level disease progression estimation.

We benchmark PILOT-GM-VAE against PILOT, PhEMD, QOT, GloScore, and baseline approaches (PILOT-GM, Pseudobulk and cellular Proportions) in ten large single cell disease atlases with up to 250 samples and 1 million cells. We demonstrate that PILOT-GM-VAE has superior results on inferring sample-level clusters or disease trajectories, recapitulating sample disease status. Moreover, we revisit and evaluate the fact that some multi-center disease atlases [covid-19 (Ren *et al*., 2021); lung (Sikkema *et al*., 2023)] have been previously shown to be affected by batch effects (Joodaki *et al*., 2024), i.e., the city of sampling or study of the samples dominated the signal of the sample-level analysis. Finally, we showcase the power of PILOT-GM-VAE in analyzing a sample-level trajectory on a breast cancer disease atlas (Kumar *et al*., 2023).

## 3 Methods

PILOT-GM-VAE receives as input a matrix *X* ∈{ℝ}^*N* ×*D*^ representing an integrated disease atlas of single cell data, where *N* represents the number of cells and *D* the dimension of the data (genes or principal components, etc.). Here, *x* corresponds to an observation (cell) from *X*. Moreover, the vector *s* = {*s*_1_,…, *s*_*N*_}, where *s*_*i*_ ∈ {1,…, *L*} indicates which sample (patient) the cells belong to.

### 3.1 Mixture of Gaussians VAE

#### 3.1.1. Generative Process

The generative model defines the joint probability of the observed data *x*, the latent variable *z*, and a discrete variable *y*, which describes the component assignments:

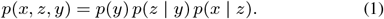

where

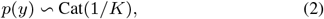

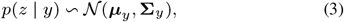

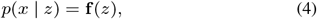

where *K* is the number of mixture components, ***µ***_*y*_ and Σ_*y*_ are the mean and covariance of the latent variable *z* conditioned on *y*, and **f**(*z*) is a neural network decoder that maps the latent variable *z* to the reconstructed data x′.

#### 3.1.2. Inference Process

The posterior distribution of the latent variable and component assignment is approximated as:

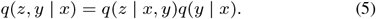

where *q*(*y* | *x*) is modeled using a categorical distribution parameterized by an encoder neural network with a Gumbel-Softmax layer as in Jang *et al*. (2016); and *q*(*z* | x, *y*) is modeled as a Gaussian distribution with parameters ***µ***_***y***_ and Σ_***y***_ produced by the encoder network. To sample from this distribution, we use the VAE re-parameterization trick (Kingma *et al*., 2014):

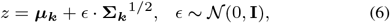

### 3.2. Objective Function

The VAE-GM optimizes a weighted Evidence Lower Bound (ELBO) composed of three terms as follows:

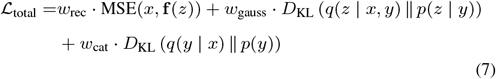

where the first term is the mean square error between the input *x* and the reconstructed output *x*_rec_, the second and third terms represent the Kullback-Leibler divergence between the distributions of the latent spaces of *z* and y and the prior distributions. The weights *w*_rec_, *w*_gauss_, *w*_cat_ control the contribution of each term.

For the encoder, we used two hidden layers with 50 units each, followed by ReLU activation functions (Nair and Hinton, 2010). The decoder network reconstructs the input data *x*_rec_ from the latent variable *z* using two fully connected layers. The latent space *z* is a 50-dimensional vector. The model was trained using the Adam optimizer (Kinga *et al*., 2015) with a learning rate of 0.001. We use a training/validation split of 80% for training and 20% for validation. The batch size is set to 32, ensuring efficient and stable training. To check the convergence of the loss function, we provide an example for the kidney data set (Lake *et al*., 2021) in Supp. Fig. S1A. Also, we checked the clustering performance of distinct weight parameters in the kidney data set (Supp. Fig. S1B). We adopt the optimal value of this evaluaiton (*w*_rec_ = 2, *w*_gauss_ = 1, and *w*_cat_ = 1) for all data sets.

#### 3.2.1. Assignment of Cells to Components

After the training of the GM-VAE, we use final posterior probabilities (*q*(*y* | *x*)) to relate cells to components. These estimates are given as input for a standard GM estimation (Pedregosa *et al*., 2011) with *K* components for refinement. This is used to to define a cluster *I*_*k*_ as:

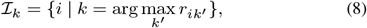

where *r*_*ik*_ is the posterior probability of cell *i* to belong to component *k*. We can similarly define a sample specific cluster 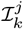 by only considering cells *i* associated with a sample *j*, i.e. *s*_*i*_ = *j*

### 3.3 Patient Specific Mixture Models and Optimal Transport

Next, we estimate parameters for a Gaussian of mixture as Θ^*j*^ = (μ^*j*^, Σ^*j*^, α^*j*^), where 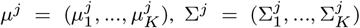, and 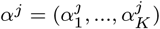, for patient *j* as follows:

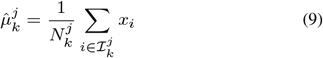

where 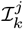 is the sets of cells associated to a component *k* and patient *j*,and 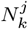 is the number of cells associated with sample *j* allocated to component *k*. Similarly, the covariance and mixing coefficients can be estimated as:

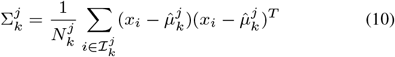

and

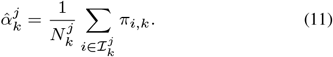

#### 3.3.1. Wasserstein Distance between Patients

To quantify the difference between the distributions of two patients (source and target) represented by a mixture of gaussian’s with parameters Θ^*s*^ and Θ^*t*^, we employ the Wasserstein Distance between two Gaussian mixtures defined as follows(Bonneel *et al*., 2011):

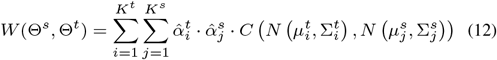

where *K* is the number of components in the Gaussian mixture model, and the cost function as the Bures-Wasserstein Distance (Bhatia *et al*., 2019) is given by:

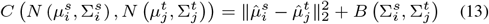

The Bures term (B) quantifies the difference in shape between two covariance matrices, which is defined as follows:

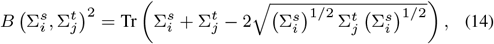

where Tr is the trace operator.

#### 3.3.2. Sample-level clustering and trajectory analysis

With the distance matrix *W* (Eq. 12), we can perform sample-level clustering by using the Leiden algorithm (Traag *et al*., 2019); or disease trajectories through diffusion maps (Coifman and Lafon, 2006). In short, diffusion maps construct a similarity matrix using a Gaussian kernel with a bandwidth function:

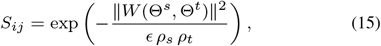

where ∈ is the scale parameter, and ρ_*s*_, ρ_*t*_ are local bandwidth functions that adapt to the density of the data (Berry and Harlim, 2016). The diffusion map embedding is derived from dominant eigenvectors (eigenvectors with the highest eigenvalues) of the normalized similarity matrix. Next, the EIPLGraph algorithm (Albergante *et al*., 2020) is used to fit a graph to the diffusion map and to infer the backbone of the trajectory. Start points of trajectories were selected by visual inspection, i.e., selection of regions related to control samples. This provides a ranking of samples based on a disease progression score *t* = {*t*_1_, *t*_2_,…, *t*_*L*_}, where *t*_*l*_ represents the rank of the sample *l*. In this study, only the two dominant eigenvectors were utilized for diffusion maps. Of note, this supervised step, i.e. selection of root cells, is only important for data interpretation and do not influence benchmarking results, which are invariant to this.

### 3.4 Datasets

We collected ten publicly available single cell disease atlases for the benchmarking of sample-level analysis methods (Table 1). These datasets were chosen based on the availability of at least two sample-level labels (control and disease). We used the same normalization, dimension reduction, and cell type annotation as in the original study unless this was not available. The cell type annotation is used as input by some of the algorithms that require it (such as PILOT, QOT, and Proportions). Eight datasets were used as previously described in Joodaki *et al*. (2024). Two of these datasets were updated to include additional or novel samples: kidney disease (Lake *et al*., 2021) and myocardial infarction (Kuppe *et al*., 2022). The novel and re-analyzed datasets are described below.

**Table 1.** Characteristics of the datasets used for benchmarking, including the total number of cells, the total number of samples, the total number of cell types reported in the original study, and the number of classes (disease status) of the samples. We also provide the disease severity scores associated with the disease status.

| Data | #Cells | #Samples | #Cell_types | #Classes & Scores |
| --- | --- | --- | --- | --- |
| Breast cancer (Kumar *et al.*, 2023) | 714,331 | 126 | 10 | 2; (Control=1, Cancer=2) |
| Covid-19 PBMC (Ren *et al.*, 2021) | 993,171 | 151 | 10 | 3; (Normal=1, Mild=2, Isch=3) |
| Diabetes (Hrovatin *et al.*, 2022) | 264,235 | 52 | 13 | 4; (Normal=1, EPD=2, TypeI&II=3) |
| Kidney (Lake *et al.*, 2021) | 104,314 | 45 | 14 | 3; (Normal=1, CKD=2, AKI=3) |
| Kidney cancer (Zvirblyte *et al.*, 2024) | 50,236 | 17 | 6 | 2; (Normal=1, Cancer=2) |
| Lupus PBMC (Perez *et al.*, 2022) | 1,263,676 | 261 | 11 | 2; (Normal=1, SLE=2) |
| Lung (Sikkema *et al.*, 2023) | 941,504 | 165 | 12 | 5; (Normal=1, COPD=2, SCLC, LA, NSCLC=3) |
| Myocardial infraction (2) (Kuppe *et al.*, 2022) | 132,888 | 23 | 11 | 3; (Normal=1, Isch=2, Fib=3) |
| Myocardial infraction (1) (Kuppe *et al.*, 2022) | 115,517 | 20 | 11 | 2; (Normal=1, IZ=2) |
| Pancreas (PDAC) (Peng *et al.*, 2019) | 57,530 | 35 | 10 | 2; (Normal=1, Cancer=2) |

#### 3.4.1. Processing of single cell sequencing data

We used single cell and single-nucleus assays from kidney samples provided by the Kidney Precision Medicine Project (Lake *et al*., 2021). Specifically, we included single cell RNA-seq from kidney tissues encompassing 104,314 cells organized into 14 major cell types. This dataset consists of samples from 45 donors: 18 controls, 12 with acute kidney failure, and 15 with chronic kidney disease. This data was obtained from CellxGene.

We now include an extended version of the myocardial infarction atlas (Kuppe *et al*., 2022), which includes 23 samples representing controls (13), ischemic (5), and fibrotic (5) samples (MI-2). The data contains 132,888 cells clustered at 11 cell types. The data was obtained from CellxGene.

The breast cancer atlas provides a detailed genomic and spatial characterization of the adult human breast. This comprehensive dataset comprises 714,331 single cells from 126 donors (Kumar *et al*., 2023), including samples from 86 normal breast tissues and 40 cancerous tissues. The cells are categorized into 10 distinct cell types. The pre-processed data was obtained from GEO (GSE195665).

The single cell transcriptional profiling of clear cell renal cell carcinoma (ccRCC) (Zvirblyte *et al*., 2024) provides a comprehensive dataset characterizing the tumor microenvironment using single cell RNA sequencing. The dataset encompasses 17 samples, including 8 normal and 9 tumor samples, and identifies 6 major cell types. The pre-processed data was obtained from GEO (GSE242299).

### 3.5 Competing methods

We describe here the evaluated competing approaches. By default, we provided the same input data for all methods. Some methods require cell types such as QOT, PILOT, and Proportions, which were defined as explained above. Other approaches have internal clustering or mixture model estimates. These methods were executed to return the same number of cluster/components as the number of cell types described in the data (Table 1). All methods provided results as a sample-level distance matrix, which was used as input for the same clustering (Leiden) and trajectory analysis (diffusion maps followed by EIPLGraph pseudotime inference) pipeline described above.

**PhEMD** processes single cell data including clustering of cells with the Monocle pipeline (Trapnell *et al*., 2014). For all datasets, we executed PhEMD with default parameters. An exception is the distribution models, which were chosen per dataset in order to optimize the clustering results^3^. For breast cancer, kidney cancer, kidney, MI (2), covid, lupus, and diabetes, we selected the Gaussian distribution; for lung, we used the Tobit distribution, and the negative binomial was used for the remaining datasets.

**Quantized Optimal Transport (QOT)** (Wang *et al*., 2025) estimates Gaussian Mixture Models for every cluster in integrated single cell data and uses the estimated parameters to find distances using optimal transport. We provided integrated, normalized, and cell-labeled scRNA-seq data as input and ran QOT (exact method) with standard parameters. By default, two components were estimated per cell type. As an exception, for MI(1), MI(2), and kidney datasets, the number of components was set to one per cluster, due to failures when running QOT with the default parameter.

**GloScope** is another sample-level approach, which uses GMMs to model sample-specific single cell data and measures distances using the KL divergence. Here, we set the number of components in GMMs to correspond to the number of cell types in each dataset.

**PILOT** is a parameter-free approach, but requires cell type annotations for estimation of sample-level distributions (Joodaki *et al*., 2024).

We also implemented several baseline approaches. The first one builds **Pseudo-bulk** libraries by summing gene expression counts for all cells. The Poisson distance Witten (2011) is then used to calculate differences between samples based on these aggregated counts. Another baseline approach is to determine the **Proportions** of each cell type within each sample from the single cell data, using the cosine distance between samples. A final baseline approach here is to use the Gaussian Mixture estimation algorithm from scikitlearn (Pedregosa *et al*., 2011) on the scRNA-seq data (at the sample-level) and use the OT approach proposed before (Eq. 12) to detect distances. This is equivalent to skipping the GM-VAE part of PILOT-GM-VAE and is denoted as **PILOT-GM**. Notably, for a fair comparison, the number of components in PILOT-GM, PILOT-GM-VAE, QOT, and GloScope is set to the number of cell types in the data.

## 4 Results

### 4.1 Benchmarking of Patient-Level Clusters and Trajectories

We applied PILOT-GM-VAE and eight competing methods [QOT (Wang *et al*., 2025); GloScope (Wang *et al*., 2024); PhEMD (Chen *et al*., 2020)); PILOT (Joodaki *et al*., 2024)] and baseline methods (PILOT-GM, Pseudobulk, Proportions, and PhEMD) to ten publicly available disease cell atlas with varying numbers of cells, samples and disease entities (Table 1). First, we evaluated the accuracy of methods in a clustering task. For this, we provided all estimated distance matrices to the Leiden algorithm (Traag *et al*., 2019), where the resolution was set to identify the corresponding number of classes (sample labels) in the data; and we contrasted the clustering labels with the sample labels using the adjusted Rand Index (Hubert and Arabie, 1985), and used the Friedman-Nemenyi (Nemenyi, 1963) test to evaluate the ranking of methods over all datasets. Regarding clustering accuracy (Fig. 2.A), PILOT-GM-VAE has the highest mean ARI value among all competing approaches. The FN test indicates that PILOT-GM-VAE’s ranking is significantly higher (p-value *<* 0.05) than PhEMD, Proportions, and GloScope methods (see Supp. Table S1 for complete results).

**Fig. 2:**
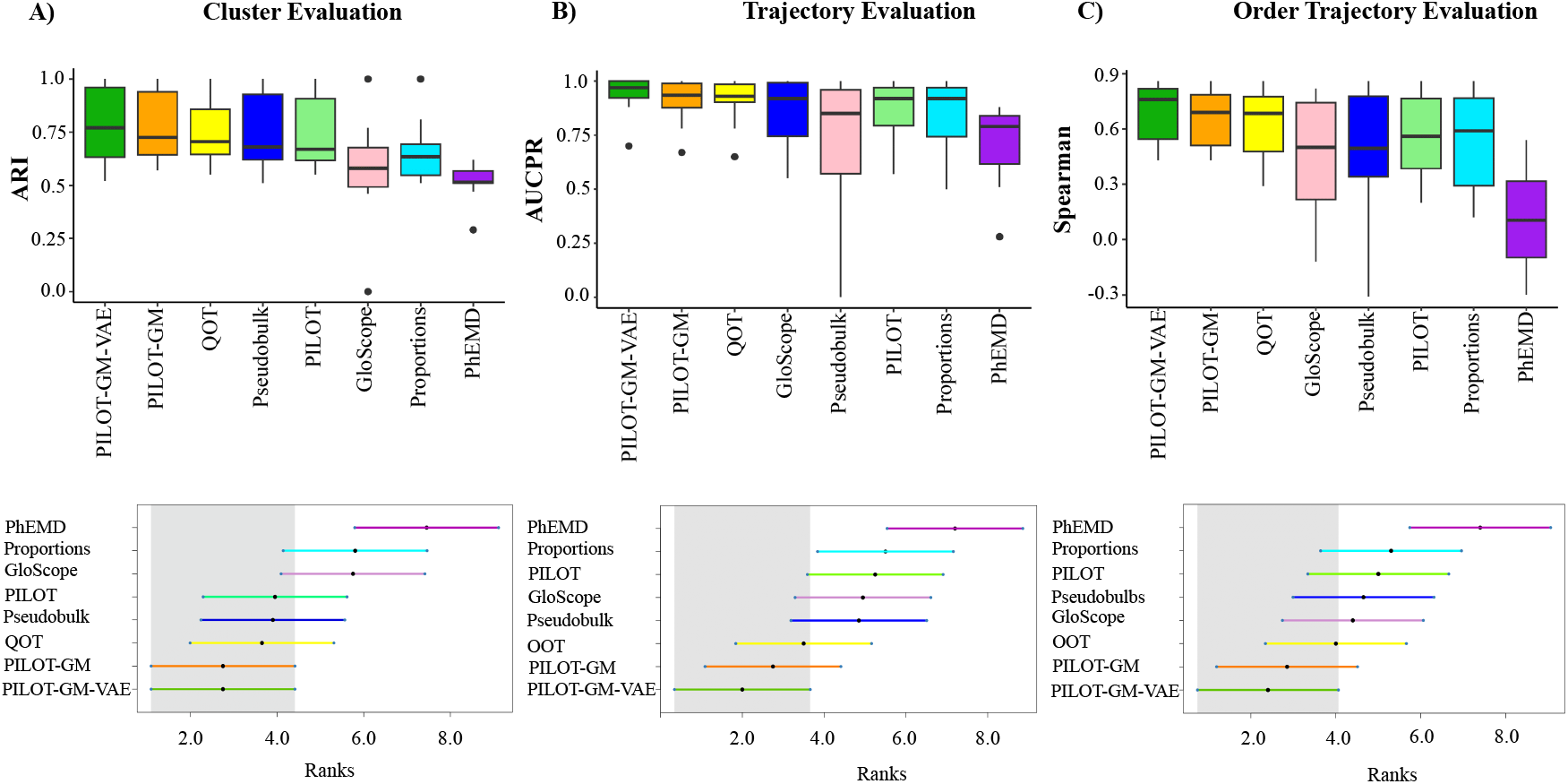
(A) Box plots showing clustering evaluation based on the Adjusted Rand Index (ARI) across methods. The x-axis represents the methods, and the y-axis represents the ARI score indicating the accuracy of patient-level clustering. Below the box plots, we show the ranking of methods (x-axis; lower values indicate best performer) in accordance with the Friedman–Nemenyi test ranks. The gray area represents the confidence interval (p-value < 0.05) of the top-performing method (lowest rank value). (B) and (C) similar to (A) for the AUCPR and Spearman correlation analysis in the evaluation of the trajectory inference problem.

Another evaluated task is the ability to infer disease progression trajectories. For this, we use the sample-level distances as input to a diffusion map algorithm (Coifman and Lafon, 2006) followed by a trajectory and pseudotime inference (Albergante *et al*., 2020). First, we evaluate whether the trajectories produced by the algorithms (patient-level pseudotimes) discriminate the sample classes (disease vs. non-disease) using the area under the precision and recall curve (AUCPR). As seen in Fig. 2B, PILOT-GM-VAE has the highest AUCPR than all competing approaches. The FN test indicates that PILOT-GM-VAE significantly outperforms PhEMD, Proportions and PILOT. Examples of diffusion maps and trajectories for PILOT-GM-VAE, PILOT-GM, QOT and Pseudo-bulk can be seen at Supp. Fig. S2.

Finally, for multi-class datasets, we can derive a disease severity score (last column of Table 1). For example, in the covid-19 dataset (Ren *et al*., 2021), we score controls as 1, mild infections as 2, and severe infections as 3. We estimate the Spearman Correlation between the pseudotime and disease severity score. As seen in Fig. 2C, PILOT-GM-VAE has the highest Spearman correlation among all competing approaches. The FN test indicates that PILOT-GM-VAE ranking is significantly higher (p-value *<* 0.05) than PhEMD. Altogether, these results collectively underscore the advantage of PILOT-GM-VAE in accurately capturing patient-level clusters and complex trajectories compared to other state-of-the-art approaches.

### 4.2 Benchmarking: computation resources

Another important aspect is the computing resources required for the methods. Here, we focus on the top four methods of trajectory benchmarking: QOT, GloScope, PILOT-GM-VAE, and PILOT-GM. Note that other approaches (PILOT, Pseudo-bulk, and Proportions) require fewer computational resources, as they are based on simpler representations of the data, however, these methods were mostly outperformed in the previously described cluster and trajectory accuracy benchmarking.

Altogether, PILOT-GM was the fastest approach followed by QOT, PILOT-GM-VAE and GloScope. Regarding memory, GloScope had the lowest requirement, while QOT, PILOT-GM, and PILOT-GM-VAE had overall similar requirements (Supp. Fig. S3 and Tables S2-S3). In the largest dataset, lupus PBMC with 1 million cells and 261 samples, PILOT-GM-VAE performed the analysis in less than seven hours, while requiring 41GB of memory. These results indicate that PILOT-GM-VAE (and the competing approaches) can be used to analyze complex disease single cell atlas data with millions of cells on a high-end notebook or desktop computer.

### 4.3 Effect of Batches in Sample-Level Analysis

A previous study on sample-level analysis of disease single cell atlases (Joodaki *et al*., 2024) indicated that the batch effect had a high impact on overall results. This is particularly true for large disease atlases such as the lung (Sikkema *et al*., 2023), breast cancer (Kumar *et al*., 2023), and covid-19 (Ren *et al*., 2021), which were based on multi-center and/or meta-analysis of distinct scRNA-seq studies. To investigate the effect of batch effects on our benchmarking analysis, we estimated the association of the inferred trajectory and sample covariates such as the location of the sample collection, study of origin, and demographic information using an ANOVA F-statistic metric. Here, we consider that methods less affected by batch effects should have a higher F-statistic for the class label than any other covariate of the study. Our results (Fig. 3) show that PILOT-GM-VAE is the only approach, which ranks disease status as the most significant covariate in all three multi-center datasets. On the other hand, the QOT prediction for covid-19 data was associated with the city of sample collection, and PILOT-GM results were mostly related to the sample source (breast cancer atlas) and study source (lung disease atlas) (Fig. 3). These results support that PILOT-GM-VAE predictions are less influenced by potential batch artifacts than competing methods.

**Fig. 3:**
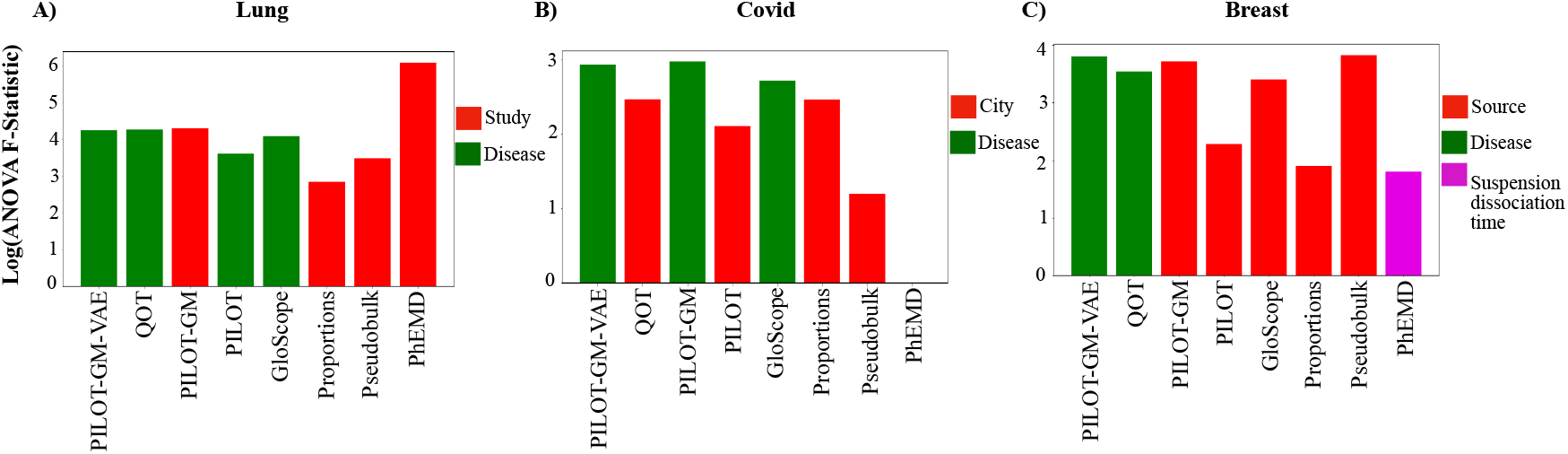
(A) Log-transformed ANOVA F-Statistic (y-axis) for the covariate with maximum F-statistic against evaluated methods (x-axis) in the lung scRNA-seq atlas. The bar color indicates whether the inferred trajectory is mostly associated with the disease label (green) or with covariates associated with batch effects (red or pink). Similar plots for (B) covid and (C) breast cancer disease atlas.

### 4.4 Disease progression in breast cancer disease atlas

To exemplify the power of PILOT-GM-VAE in interpretation, we apply PILOT-GM-VAE to a breast cancer disease atlas (Kumar *et al*., 2023). Trajectory analysis can discriminate the majority of controls from cancer samples (Fig. 4A-B). For this, PILOT-GM-VAE used mixture models with ten components. Out of these, seven components corresponded directly to one of the ten major cell types described in Kumar *et al*. (2023) (Supp.Fig. S4). Among other differences, PILOT-GM-VAE found two components associated with fibroblast cells (components three and six), which we denote here as Fibroblast 1 and 2. Moreover, these subpopulations were not related to fibroblast subtypes described in Kumar *et al*. (2023) (Supp. Fig. S4C).

**Fig. 4:**
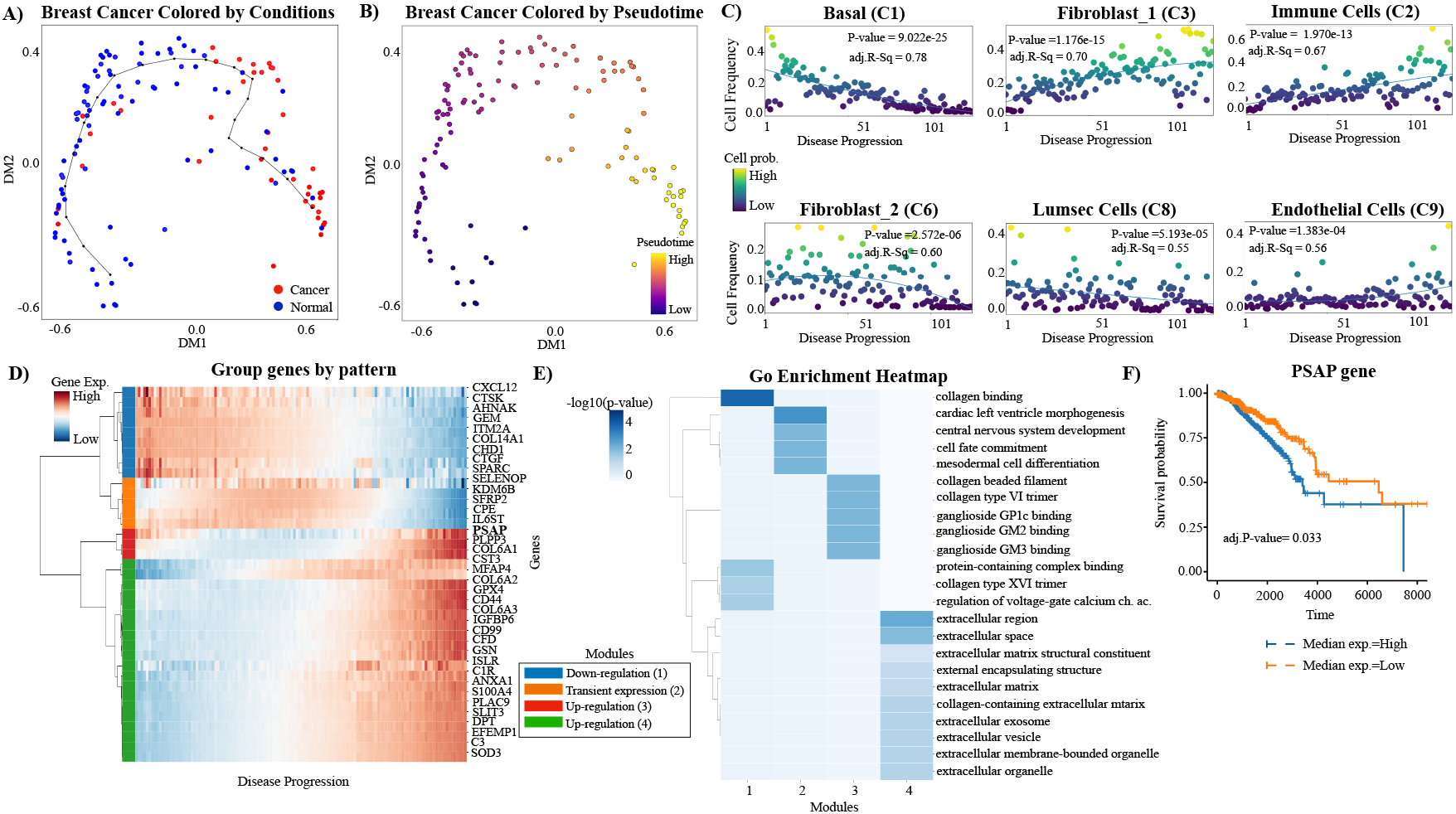
(A) Diffusion map of the breast cancer disease atlas. The line represents the predicted disease trajectory. (B) Diffusion map of the breast cancer disease atlas, colored by pseudotime (disease progression). (C) Proportion (y-axis) of the top six components (annotated) (p-value < 0.05 and highest R2) vs. disease progression (x-axis). (D) Gene modules detected in genes with expression specific to Fibroblast 1 (C3) and with significant changes across disease progression (x-axis). (E) GO enrichment of the genes for each cluster. (F) Kaplan-Meier plot showing survival of TCGA breast cancer patients stratified by median expression of the PSAP gene. p-values were estimated based on the top 15 genes (log-rank test) and corrected with the Benjamini-Hochberg procedure.

One interesting downstream analysis is to check for how the proportion of cells associated with particular components changes over the inferred disease trajectory by the use of non-linear regression models (Huber, 1965). This detected six components, in which the proportion of cells are related to the inference disease trajectories (p-value < 0.0001; Fig. 4C). This indicated a loss of epithelial cells (basal C1 and lumsec cells C8) and an increase in immune cells (C2) and endothelial cells (C9). Interestingly the two components related to fibroblast cells (C3 and C6) have distinct changes in cell trajectories, i.e., while Fibroblast 1 (C3) increases along the disease trajectory, Fibroblast 2 (C6) decreases, particularly for samples at the end of the trajectory.

To gain further insights into the mechanisms associated with the cancer-associated Fibroblast 1 (C3), we performed a non-linear regression test (p-value < 0.05) to find genes with changes associated with the disease trajectory. This was followed by a Wald test (Van den Berge *et al*., 2020) (p-value < 0.01, FC > 0.5) to find genes whose expression is specific to Fibroblast 1 (C3) cells (Joodaki *et al*., 2024). A clustering analysis of these genes reveals four main gene modules (1- down-regulation, 2 - transient expression, and 3 and 4 up-regulation). Functional analysis (GO enrichment (Reimand *et al*., 2007)) indicates that genes with transient expression (module 2) are related to cell differentiation, and clusters associated with up-regulation (modules 3-4) are related to collagen protein and the extracellular matrix (Fig. 4E). This indicates that these cells undergo cellular differentiation, leading to fibrosis-associated expression programs.

Fibrosis is supportive of the tumor environment and related to poor prognosis in cancers (Piersma *et al*., 2020), including breast cancer (Quintela-Fandino *et al*., 2024). We next performed survival analysis using gene expression data from the TCGA breast cancer database (13 *et al*., 2012) among the top 15 genes related to Fibroblast 1 cluster, we found that the PSAP gene was associated with an increase in mortality (Fig. 4F). This gene is part of gene module 3 (Fig. 4D) and shows increased expression in disease progression. This gene has been previously related to the activation of androgen and the promotion of breast cancer growth (Ali *et al*., 2015; Wu *et al*., 2012). Altogether, this analysis highlights how PILOT-GM-VAE predicted trajectories can be used to highlight cellular and molecular changes in a disease single cell atlas.

## 5 Conclusion

In this manuscript, we describe PILOT-GM-VAE, which integrates a Gaussian Mixture Variational Autoencoder with optimal transport for patient-level clustering and trajectory analysis. The use of Gaussian mixtures was previously explored by GloScope (Wang *et al*., 2024). However, GloScope uses a Kullback–Leibler (KL) distance to contrast GM distributions. In contrast to the Wasserstein Distance, KL is not a metric and ignores the expression information of cells. While QOT (Wang *et al*., 2025) also used a similar Wasserstein Distance as PILOT-GM-VAE, it estimates mixtures based on previous cell clustering. Since single cell clustering are mostly based on non-parametric graph-based algorithms such as Leiden (Traag *et al*., 2019), it is possible that this leads to poor GM fits. Moreover, the use of VAE-GMs indicated better results in contrast to EM estimated GMs (see PILOT-GM vs PILOT-GM-VAE). This highlights the effectiveness of GM-VAE in modeling complex and noisy single cell data.

Our evaluation also included the influence of batch-related covariates in the sample-level analysis, an issue that is largely overlooked in the literature. Once again, PILOT-GM-VAE was the only approach not directly influenced by variables such as the city of sampling or the study origin. A final aspect of our benchmarking is the computational requirements, which indicates higher computational costs for more complex approaches as PILOT-GM-VAE and GloScope. However, all methods can be executed in less than a day with a high-end desktop computer, which is a minimal effort compared to the generation of the disease cell atlas. Altogether, these results highlight the advantage of PILOT-GM-VAE in sample-level analysis tasks such as clustering and trajectory analysis.

Another previously poorly explored is the interpretability of sample-level analysis results. While previous approaches, such as PILOT and QOT, rely on previous clustering (or cell type annotation) to facilitate the interpretation of cell type compositional changes, the use of mixtures requires additional annotation of their clusters (or component associations). In a case study of the breast cancer disease atlas. We showed that clustering based on the GM aligned well with the major cell types reported in the original study. Interestingly, one exception supported the sub-clustering of fibroblasts, where compositional changes were inverted in the disease trajectory. Functional analysis indicated an increase in a fibrosis-related fibroblast population with disease progression. Moreover, genes with expression changes related to Fibroblast 1 cells were prognostic of disease outcome. This further supports the power of PILOT-GM-VAE in uncovering novel biological insights in into progression.

Some open/future research questions are the extension of PILOT-GM-VAE in spatial disease atlas; or multimodal cell atlas. The first would require the definition of spatially-aware Gaussian mixture estimation approaches. The latter problem could be answered with kernel fusion approaches. Nevertheless, both multimodal and spatial disease atlas data are scarce, which currently impairs a comprehensive benchmarking of computational approaches.

## Acknowledgments

This work was funded by grants of the Deutsche Forschungsgemeinschaft KFO5011 (Projekt ID: 445703531) and by the Bundesministerium für Bildung und Forschung (BMBF e:Med Consortia Fibromap and Graphs4Patients).

## Source Code and Data Availability

The software, code, trained models, estimated distances for all datasets, and tutorials are available at https://github.com/CostaLab/PILOT-GM-VAE/tree/main. All pre-processed and annotated data is available as python AnnData objects in https://zenodo.org/records/7956950, https://zenodo.org/records/7957118 and https://zenodo.org/records/14615923.

## Footnotes

1 Note that a disease single cell atlas cannot be defined as a 3-dimensional tensor, as single cell dimensions cannot be trivially mapped across samples.

2 The original method from Kingma was based on a semi-supervised problem.

3 This procedure made PhEMD results over-optimistic.

## Notes

https://github.com/CostaLab/PILOT-GM-VAE/tree/main

https://zenodo.org/records/7956950

https://zenodo.org/records/7957118

https://zenodo.org/records/14615923

### Competing Interest Statement

The authors have declared no competing interest.

