## Supplementary material for "PILOT-GM-VAE: Patient-Level Analysis of single cell Disease Atlas with Optimal Transport of Gaussian Mixture Variational Autoencoders": PILOT_GM_VAE_Supp.pdf

**Supplementary Materials**

Table S1. Benchmarking of area under the PR curve (AUCPR) and clustering results (ARI). Results in bold show the best value per method.

| Methods | Metrics/Datasets | Breast | Covid | Diabetes | Kidney cancer | Kidney | Lupus | Lung | MI_1 | MI_2 | PDAC |
|---|---|---|---|---|---|---|---|---|---|---|---|
| **Pseudobulk** | ARI | 0.62 | 0.51 | 0.67 | **1.00** | 0.62 | 0.57 | 0.69 | **1.00** | 0.71 | **1.00** |
| | Spearman | 0.34 | 0.16 | 0.34 | **0.86** | 0.37 | 0.62 | -0.31 | **0.83** | 0.71 | **0.80** |
| | AUCPR | 0.55 | 0.92 | 0.74 | **1.00** | 0.76 | 0.92 | 0.59 | **1.00** | 0.97 | **1.00** |
| **QOT** | ARI | **0.63** | 0.55 | 0.73 | **1.00** | 0.72 | 0.55 | 0.69 | 0.90 | 0.69 | **1.00** |
| | Spearman | 0.44 | 0.29 | 0.43 | **0.86** | 0.69 | 0.59 | 0.70 | 0.80 | 0.68 | **0.80** |
| | AUCPR | 0.65 | 0.94 | 0.78 | **1.00** | 0.90 | 0.91 | 0.92 | 0.82 | 0.94 | **1.00** |
| **PILOT-GM-VAE** | ARI | 0.55 | 0.52 | **0.77** | **1.00** | **0.84** | 0.61 | **0.70** | **1.00** | **0.77** | **1.00** |
| | Spearman | **0.51** | **0.43** | 0.48 | **0.86** | **0.84** | 0.65 | 0.72 | 0.82 | **0.81** | **0.80** |
| | AUCPR | **0.70** | **0.96** | **0.88** | **1.00** | **1.00** | **0.93** | 0.92 | **1.00** | **0.98** | **1.00** |
| **GloScope** | ARI | 0.53 | 0.46 | 0.67 | 0.48 | 0.61 | null | 0.68 | 0.55 | **0.77** | **1.00** |
| | Spearman | 0.41 | 0.39 | **0.59** | 0.16 | -0.12 | null | 0.72 | 0.82 | 0.75 | **0.80** |
| | AUCPR | 0.64 | 0.93 | 0.79 | 0.55 | 0.54 | null | 0.91 | **1.00** | 0.97 | **1.00** |
| **PILOT-GM** | ARI | 0.58 | **0.57** | 0.76 | **1.00** | 0.75 | 0.63 | **0.70** | **1.00** | 0.68 | **1.00** |
| | Spearman | 0.46 | **0.43** | 0.45 | **0.86** | 0.68 | 0.66 | **0.74** | 0.82 | 0.70 | **0.80** |
| | AUCPR | 0.67 | **0.96** | 0.78 | **1.00** | 0.86 | **0.93** | **0.93** | **1.00** | 0.94 | **1.00** |
| **PILOT** | ARI | **0.63** | 0.55 | 0.67 | **1.00** | 0.61 | 0.61 | 0.67 | 0.98 | 0.69 | **1.00** |
| | Spearman | 0.20 | 0.26 | **0.55** | **0.86** | 0.33 | 0.55 | 0.57 | 0.81 | 0.66 | **0.80** |
| | AUC. | 0.57 | 0.94 | 0.71 | **1.00** | 0.76 | 0.91 | 0.90 | 0.98 | 0.93 | **1.00** |
| **PhEMD** | ARI | 0.52 | 0.51 | 0.62 | 0.61 | 0.58 | 0.51 | 0.29 | 0.47 | 0.53 | 0.51 |
| | Spearman | -0.10 | -0.09 | -0.30 | 0.43 | 0.25 | 0.11 | 0.54 | -0.15 | 0.10 | 0.34 |
| | AUCPR | 0.28 | 0.88 | 0.51 | 0.81 | 0.77 | 0.58 | 0.85 | 0.81 | 0.73 | 0.87 |
| **Proportions** | ARI | 0.52 | 0.54 | 0.64 | **1.00** | 0.57 | **0.70** | 0.67 | 0.81 | 0.63 | 0.51 |
| | Spearman | 0.12 | 0.24 | 0.45 | **0.86** | 0.20 | **0.66** | 0.52 | 0.81 | 0.67 | **0.80** |
| | AUCPR | 0.62 | 0.94 | 0.50 | **1.00** | 0.69 | 0.91 | 0.90 | 0.98 | 0.93 | **1.00** |

Table S2. Comparison of computational time of selected methods in minutes. Results in bold show the lowest time per method. Results for GloScope in Lupus data are missing due to the failed execution.

| **Data/Methods** | **QOT** | **GloScope** | **PILOT-GM** | **PILOT-GM-VAE** |
|---|---|---|---|---|
| | **Time (Min)** | **Time (Min)** | **Time (Min)** | **Time (Min)** |
| Breast cancer | 37.83 | 134.41 | **37.23** | 90.48 |
| Covid-19 PBMC | 125.01 | 297.38 | **63.61** | 131.16 |
| Diabetes | 28.26 | 55.31 | **3.05** | 23.18 |
| Kidney | 3.31 | 38.5 | **2.00** | 10.21 |
| Kidney cancer | 0.93 | 26.71 | **0.51** | 7.06 |
| Lupus PBMC | 511.10 | Null | **373.78** | 408.73 |
| Lung | 213.05 | 305.9 | **79.3** | 165.45 |
| Myocardial infraction (2) | **0.80** | 18.23 | 0.81 | 13.70 |
| Myocardial infraction (1) | **0.65** | 15.16 | 0.86 | 14.56 |
| Pancreas (PDAC) | 5.81 | 21.81 | **0.80** | 8.33 |

Table S3. Comparison of memory usage (in GB) between selected methods. Results in bold show the lowest memory per method

| **Data/Methods** | **QOT** | **GloScope** | **PILOT-GM** | **PILOT-GM-VAE** |
|---|---|---|---|---|
| | **Memory (GB)** | **Memory (GB)** | **Memory (GB)** | **Memory (GB)** |
| Breast cancer | 28.65 | **0.72** | 30.39 | 30.17 |
| Covid-19 PBMC | 30.16 | **1.05** | 32.95 | 32.54 |
| Diabetes | 6.91 | **0.59** | 7.95 | 7.92 |
| Kidney | 12.02 | **0.50** | 12.08 | 12.11 |
| Kidney cancer | 7.96 | **0.47** | 8.49 | 8.21 |
| Lupus PBMC | **15.39** | Null | 18.74 | 19.07 |
| Lung | 39.25 | **1.04** | 41.95 | 41.86 |
| Myocardial infraction (2) | 11.10 | **0.49** | 5.14 | 5.02 |
| Myocardial infraction (1) | 5.90 | **0.47** | 6.38 | 6.24 |
| Pancreas (PDAC) | 3.21 | **0.46** | 3.69 | 3.37 |

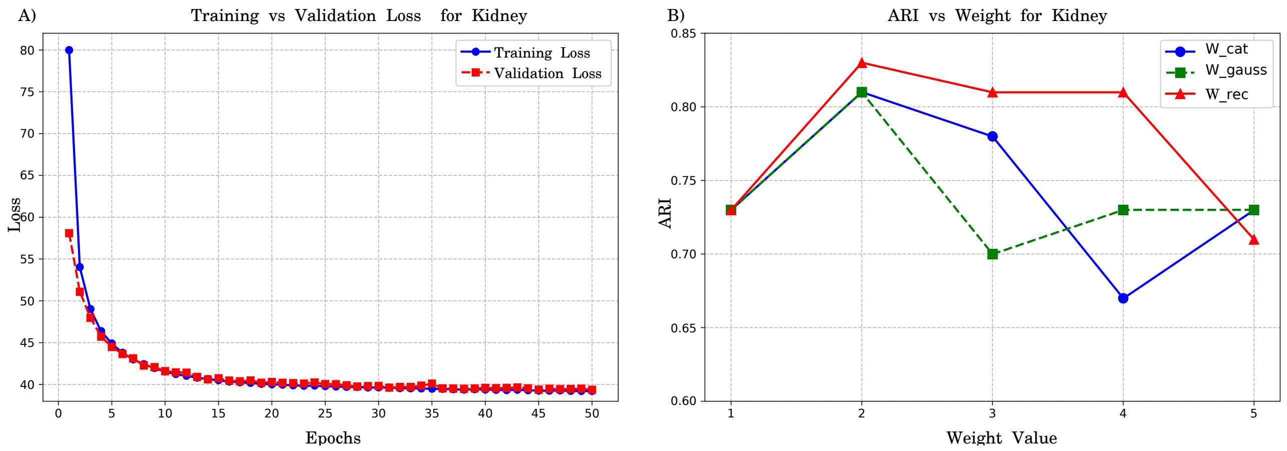


Fig. S1: (A) Training and validation loss curves over 50 epochs for the Kidney dataset. The consistent decrease in both losses indicates effective learning. (B) Line plots showing ARI values (y-axis) for distinct weight values (x-axis) in the Kidney data set. We tested weight values ranging from 1 to 5 for each of the three terms of Eq. 7 ($w_{\text{rec}}$, $w_{\text{gauss}}$, and $w_{\text{cat}}$) while keeping other weights fixed as 1.

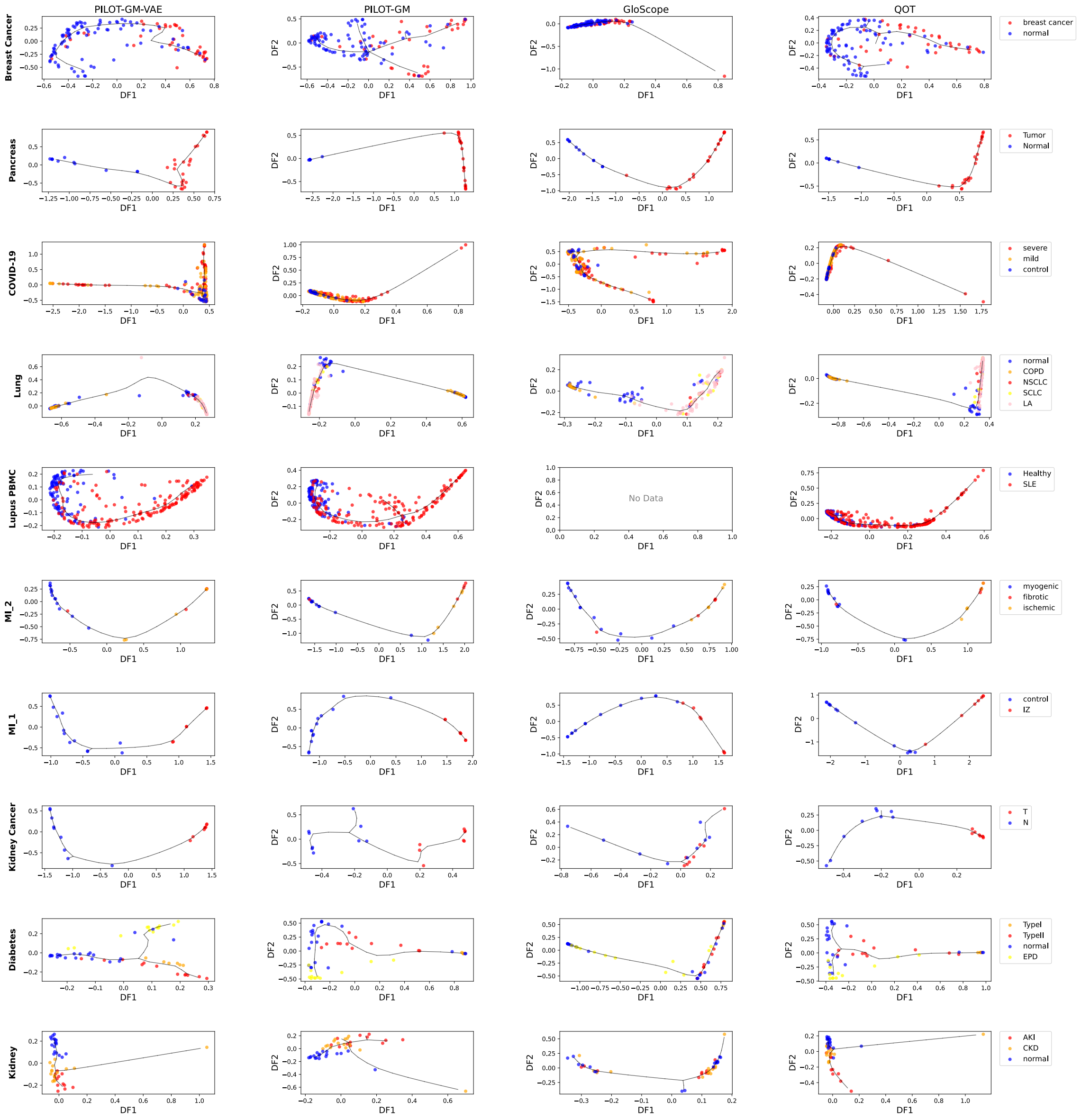


Fig. S2: Diffusion maps and pseudo-time inference for four top methods scoring methods in our benchmarking. Each point corresponds to a sample, with colors representing their disease label. The backbone trajectory is computed with ElpiGraph; and used for pseudo-time (disease progression score) inference.

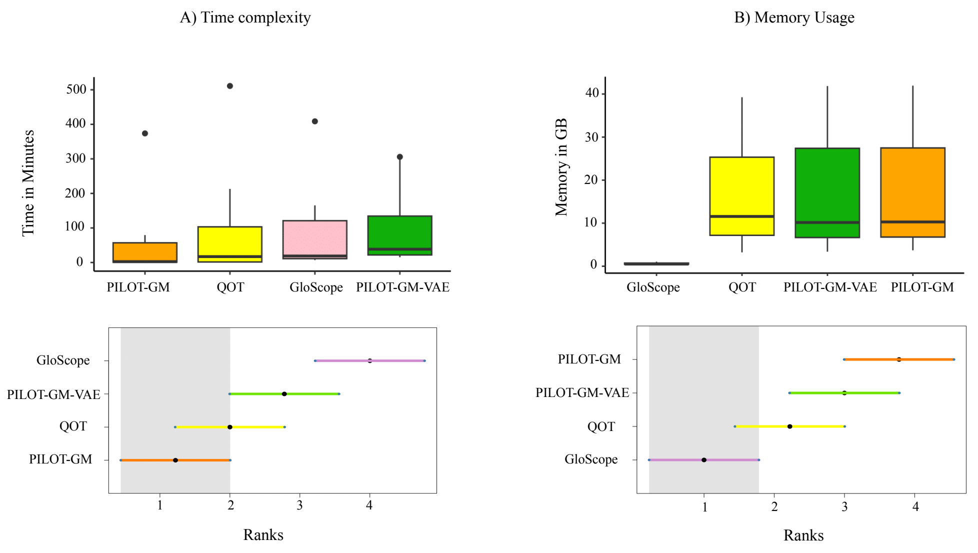


Fig. S3: (A) Box plots showing the time requirements for all datasets for top-scoring algorithms. We show below the ranking of methods (x-axis; lowest means lowest time) in accordance with the Friedman–Nemenyi test ranks. The gray area indicates the confidence interval (p-value > 0.05) of the fastest method (lowest rank). (B) is similar to (A) for the memory usage in GB.

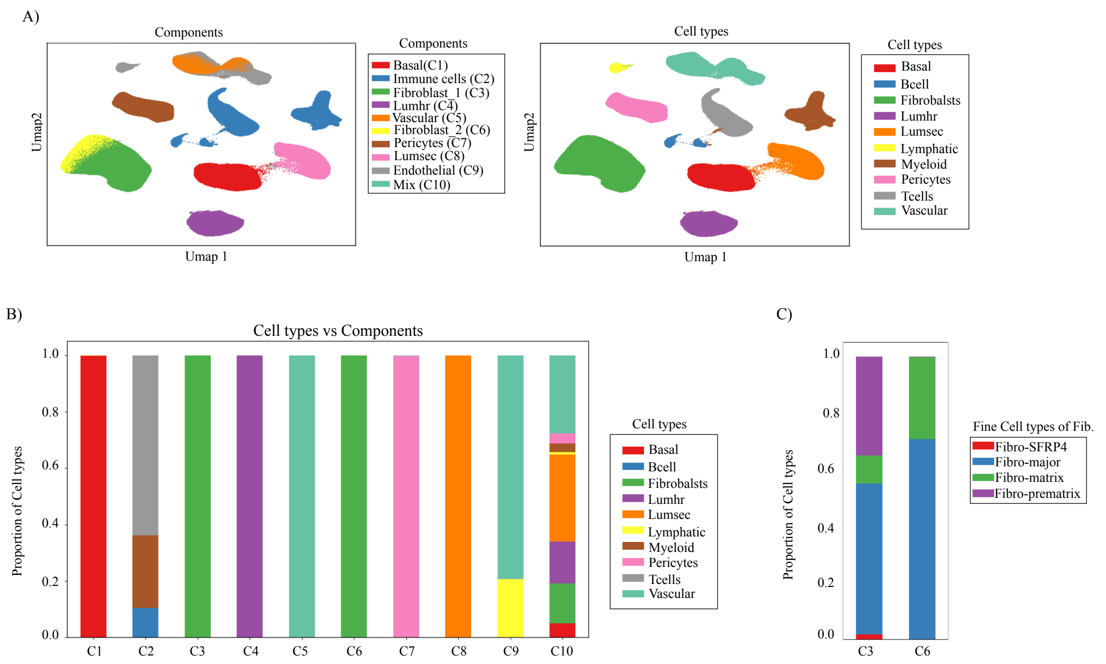


Fig. S4: (A) UMAP representation of the breast cancer disease cell atlas. On the left, you have the assignment of cells to 10 distinct GM-VAE components, and on the right, the original clustering of the cells. (B) Stacked bar plots contrasting the association of cells to components vs previously annotated cell types. C1, C4, C5, C7, C8, and C9 had a one-to-one correspondence to the original clusters from Kummar et al., The fibroblast cluster was divided into two components (C3 - Fibroblast 1 and C6 Fibroblast 2). C2 is called immune cells, as it combines B-cells, T-cells, and Myeloid cells. C5 is called endothelial as it combines both vascular cells and the closely related lymphatic endothelial cells. The component C10 was formed by a mix of cells. (C) Stacked bar plots showing the association of cells of fibroblast components C3 and C6 vs the four fibroblast subpopulations described in Kummar et al., Two of the largest clusters (Fibro-major and Fibro-matrix) were equally associated with both components, while Fibro-SFRP4 and Fibro-prematrix were only related to C3.
